# Biological sex and age shape melanoma tumour evolution

**DOI:** 10.1101/828038

**Authors:** Martin Lotz, Simon J. Furney, Amaya Virós

## Abstract

Many cancer types display sex and age disparity in incidence and outcome. Here, we establish a mathematical approach using cancer mutational data to analyze how sex and age shape the tumour genome. We model how age-related (clock-like) somatic mutations that arise during cell division, and extrinsic (environmental) mutations accumulate in cancer genomes. As a proof-of-concept, we apply our approach to melanoma, a cancer driven by cell-intrinsic age-related mutations and extrinsic ultraviolet light-induced mutations, and show these mutation types differ in magnitude, chronology and by sex in the distinct molecular melanoma subtypes.

Our model confirms age and sex are determinants of cellular mutation rate, shaping the final mutation composition. We show mathematically for the first time how, similar to non-cancer tissues, melanoma genomes reflect a decline in cell division during ageing. We find clock-like mutations strongly correlate with the acquisition of ultraviolet light-induced mutations, but critically, males present a higher number and rate of cell division-linked mutations. These data indicate the contribution of environmental damage to melanoma likely extends beyond genetic damage to affect cell division.

## Introduction

Sex and age disparity in cancer incidence and outcome are well described and studies have revealed age(Balch et al. 2013; Cavanaugh-Hussey et al. 2015) and sex differences(Gupta et al. 2015) in genomics(Li et al. 2019). Somatic mutations are drivers of cancer and mutations arise in cells due to damage following cell-intrinsic processes, as well as due to external environmental damage on the DNA strand. Recent work describes computational methods to discern the multiple, distinct signatures of DNA damage imprinted on DNA depending on the insult(Kucab et al. 2019), but to date there are no available models to study the relationship between the identified damaging processes.

We used cutaneous malignant melanoma as a model cancer to develop a mathematical framework to investigate the dependencies between cell-intrinsic, extrinsic mutational processes, sex and age. Cutaneous melanoma exemplifies a cancer type combining cell-intrinsic (cell division) and environmental (ultraviolet light) damaging processes, as well as presenting an age and sex bias. Male melanoma patients and the elderly population have a higher incidence and rate of death, so we studied if the genomic imprint of ultraviolet radiation (UVR) exposure and cell division are possible sources for the disparity.

Cutaneous melanoma presents a broad range of clinical subtypes of disease, categorized by age of onset, history and pattern of UVR exposure(Schadendorf et al. 2018; Shain and Bastian 2016). At one end of the spectrum, we identify elderly patients with melanomas arising at anatomic sites that have been chronically exposed to UVR over a lifetime with a high tumour mutation burden (TMB). In contrast, melanoma in younger patients arises decades after acute sunburns, over skin that is only intermittently exposed to UVR, with a lower TMB(Cancer Genome Atlas Network. Electronic address and Cancer Genome Atlas 2015; Shain and Bastian 2016; Whiteman et al. 2003). These broad clinico-epidemiological categories show that the relationship between melanoma incidence and UVR exposure is not linear and cumulative.

The mutually exclusive oncogenic drivers ^V600^BRAF and NRAS underpin the majority of cutaneous melanomas(Akbani et al. 2015). Loss-of-function mutations in the tumour suppressor NF1 drive an additional subset of cases, and a further subgroup is defined by the absence of ^V600^BRAF, NRAS or NF1 mutations (triple wild type; W3). These four genetically distinct categories overlap to some extent with clinical characteristics, with ^V600^BRAF being more prevalent in younger patients(Akbani et al. 2015).

The major contributors to total autosomal TMB in melanoma are mutations due to intrinsic cell processes damaging DNA or deficient DNA damage response, and extrinsic environmental processes such as UVR(Alexandrov et al. 2013). Here we examine the relationship between mutational processes and their contribution to the melanoma somatic mutation load, their variation over time and across sexes. We provide a novel approach and mathematical framework to model and integrate how the specific damage patterns in DNA arise over time and across the sexes. Analysing the strong bias in the mutational landscape could point to key biological differences in how tumours develop and evolve during ageing and across sexes.

## Results

### Clock-like and UVR-driven mutations accumulate with age at distinct rates in the molecular subtypes of melanoma

We catalogued the base substitutions in 396 whole exome cutaneous melanoma samples from TCGA (TCGA-SKM)(Akbani et al. 2015) according to 96 categories defined by the base substitution, the preceding and following bases(Alexandrov et al. 2013). 172 had a ^*V600*^BRAF (BRAF) mutation, 96 *NRAS*, 44 *NF1* and 84 samples were non-*BRAF*/non-*NRAS*/non-*NF1* (W3; Supplementary Table 1).

We inferred the mutational signatures that account for the somatic mutations from the TCGA data. We extracted the DNA mutational signature linked to UVR (Signature 7, COSMIC database), which is present to varying degrees across melanomas. Next, we identified the intrinsic, age-related signature observed in normal cells and cancers with high cell turnover, which corresponds to spontaneous deamination of methylated cytosine residues into thymine at CpG sites that remain unrepaired due to rapid DNA replication (Signature 1, COSMIC database(Alexandrov et al. 2013, 2015; Nik-Zainal et al. 2012)). This clock-like mutational process allows estimation of the number of divisions a cell has undergone since its inception. We confirmed a positive correlation between the median number of Signature 1 mutations per year and age for all samples (Spearman ρ 0.41, *P*<0.0003, Figure 1A), indicating that these mutations accumulate with age. NRAS, NF1 and W3 melanomas presented an increase in the mean Signature 1 mutations with age, but this relationship was less significant in NF1 and W3 melanomas, likely due to the lower sample size (NRAS Spearman ρ 0.35, *P*<0.01; Figure 1B). Strikingly, there was no significant rank correlation between Signature 1 and age in BRAF samples (Spearman ρ 0.03, *P*=0.41). To examine the difference in the rates of Signature 1 mutation accumulation between BRAF, NRAS, NF1 and W3 melanomas, we determined the ratio between the number of Signature 1 mutations to age, and found significant differences in the ratios of BRAF and W3 to NF1 melanomas (pairwise Wilcoxon rank-sum test with Bonferroni correction, *P*<0.0027; Figure 1C), but less pronounced for BRAF and NRAS samples.

**Figure 1.**
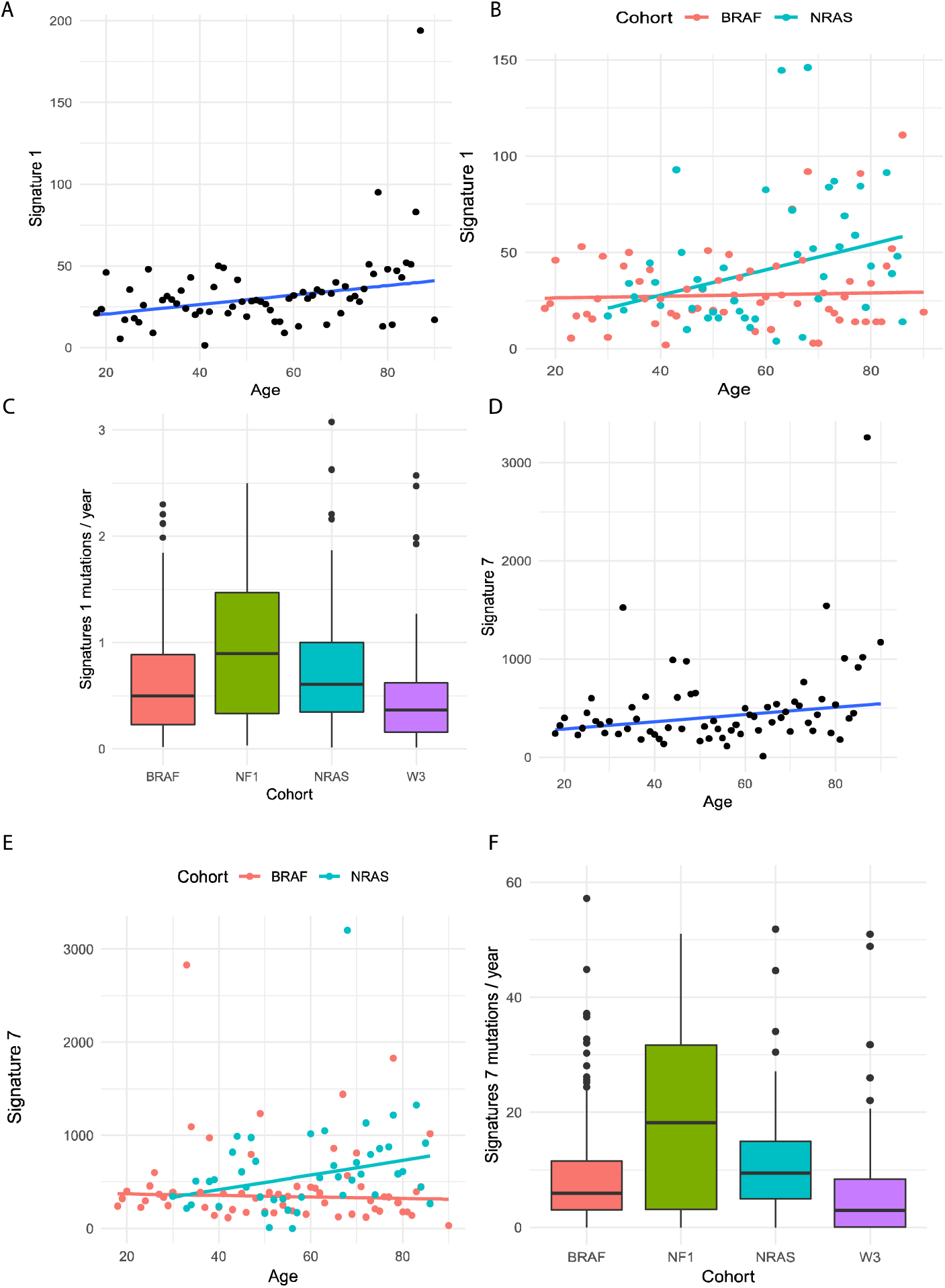
The molecular subtypes of melanoma present distinct ratios of clock-like and UVR mutations per unit of time. **(A)** Correlation analysis between somatic mutations due to clock-like Signature 1 mutations in cutaneous melanomas and age. Dots represent the median number of mutations for each age. **(B)** Correlation analysis between clock-like Signature 1 mutations in the molecular subtypes of cutaneous melanomas (BRAF: red; NRAS: blue) and age. Dots represent the median number of mutations for each age. **(C)** Ratio of number of signature 1 mutations per year across the molecular subtypes of cutaneous melanoma. **(D)** Correlation analysis between somatic mutations due to UVR Signature 7 mutations in cutaneous melanomas and age. Dots represent the median number of mutations for each age. (E) Correlation analysis between UVR Signature 7 mutations in the molecular subtypes of cutaneous melanomas (BRAF: red; NRAS: blue) and age. Dots represent the median number of mutations for each age. **(F)** Ratio of number of Signature 7 mutations per year across the molecular subtypes of cutaneous melanoma.

We next examined the contribution of Signature 7 mutations. As expected, we found a progressive increase in the numbers of mutations as patients aged, in accordance with progressive accumulation of UVR damage during the course of life (Spearman ρ = 0.37, *P*<0.006; Figure 1D). However, the rate of mutations varied depending on the molecular subtype of melanoma, with once again no significant correlation in BRAF melanomas between Signature 7 and age of diagnosis (Spearman ρ =-0.15, P=0.87, Figure 1E), and NRAS samples showing a steady increase of Signature 7 with age (Spearman ρ = 0.37, *P*<0.006). Similar to Signature 1, there was a significant difference in the ratio of Signature 7 mutations to age across the subtypes (*P*<0.03, Mann-Whitney U test with Bonferroni correction; Figure 1F).

### Ageing affects the dynamics of the mutational landscape

Common models of cancer have assumed that mutations accumulate at a linear rate over time(Stratton et al. 2009). Genetic changes accumulate from early life(Alexandrov et al. 2015; Blokzijl et al. 2016) and a decline in replicative function with age is visible in many tissues(Wang and Hekimi 2015). We used our cohorts to test the relationship between ageing and clock-like mutation rate in melanoma, and found the ratio of mutations per year decreases with age (Spearman ρ =-0.34, *P*<0.005; Figure 2A). Specifically, the decline in the number of mutations per year is pronounced in BRAF (Spearman −0.44, *P*<0.0004) but not statistically significant in NRAS or W3 melanomas. We did not include NF1 samples in the analysis, as this subtype is almost exclusive to the elderly. These data lend further support to a distinct behavior of melanocytes and melanoma cells driven by BRAF.

**Figure 2.**
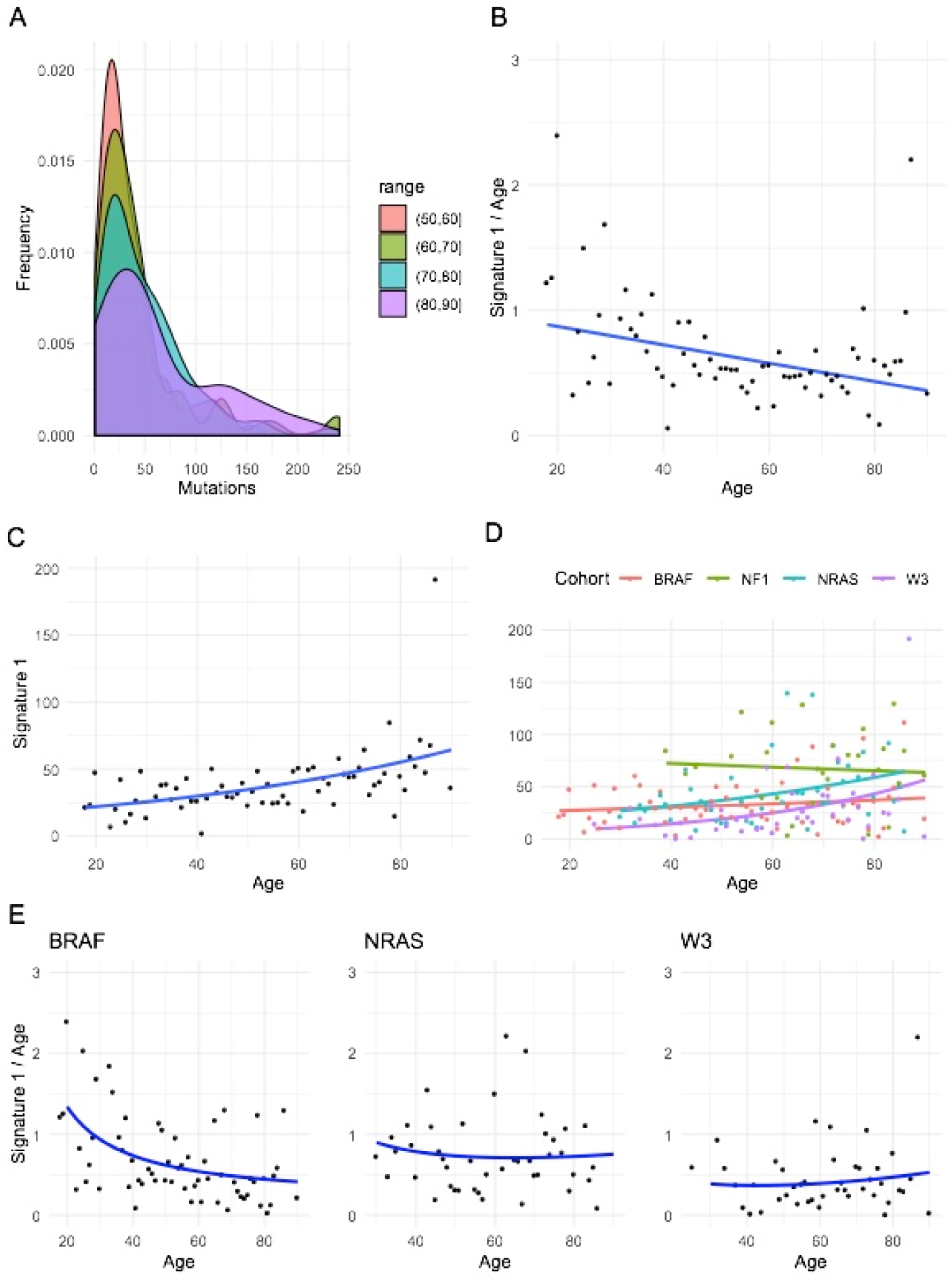
Ageing affects the intrinsic mutation rate of the molecular subtypes. **(A)** Correlation analysis between clock-like Signature 1 mutations per year and age in cutaneous melanomas. Dots represent the median number of mutations for each age. **(B)** Distribution curves displaying Signature 1 mutation frequency across age ranges (red: 50-60 year-old range; green: 60-70 year-old range; blue: 70-80 year-old range; purple: 80-90 year-old range). **(C)** Exponential model for the accumulation of Signature 1 mutations in all melanomas. This curve models the Poisson mean distribution of mutations at each age, with age-dependent rate. **(D)** Exponential model for the accumulation of Signature 1 mutations in the molecular subtypes of cutaneous melanoma (BRAF: red; NF1: green; NRAS: blue; W3: purple). This curve models the Poisson mean distribution of mutations at each age, with age-dependent rate. **(E)** Change in Signature 1 mutations per year with age across BRAF, NRAS and W3 subtypes.

To analyse the differences in ageing dynamics, we considered the mean number of mutations at each age, modelled by a Poisson distribution with age-dependent rate, and show the ratio of mutations by age decreases in older age groups (Figure 2B). We used an overdispersed Poisson (negative binomial) regression to estimate the parameters of the exponential model for each subtype BRAF, NRAS and W3; and found that overall, in this model, the amount of Signature 1 mutations increases by a multiplicative factor of e^α^ = 1.0124 per year (Figure 2C). In contrast, the increase factor is only 1.005 for BRAF, 1.0156 for NRAS and 1.0235 for W3 melanomas (Figure 2D, Supplementary Table 2). Thus, whilst the ratio of Signature 1 damage acquisition to age in BRAF and NRAS melanomas decreases during ageing, likely reflecting a deceleration of the cell proliferation rate during maturity, the ratio per year of clock mutations in W3 melanomas slightly increases during the human lifespan, reflecting a distinct behaviour (Figure 2E).

### The rate of clock-like mutations is linked to UVR damage

Previous experiments show acute UVR drives melanocyte proliferation(Viros et al. 2014), but the long-lasting effects of UVR on cell division have not been explored. We investigated the proportion of Signature 1 and Signature 7 mutations across the melanoma subtypes and found that UVR underpins approximately 75% of all mutations in BRAF, NRAS and NF1 samples, while only half of the mutations in W3 samples are accounted for by UVR (Figure 3A). Furthermore, we find a greater proportion of the ageing signature that is uncoupled from cellular division (Signature 5, characterised by T:A to C:G transitions) contributing to the overall mutational burden of W3 melanomas. The underlying biological process driving Signature 5 mutations is unknown, but is linked to ageing independently of cellular division(Blokzijl et al. 2016).

**Figure 3.**
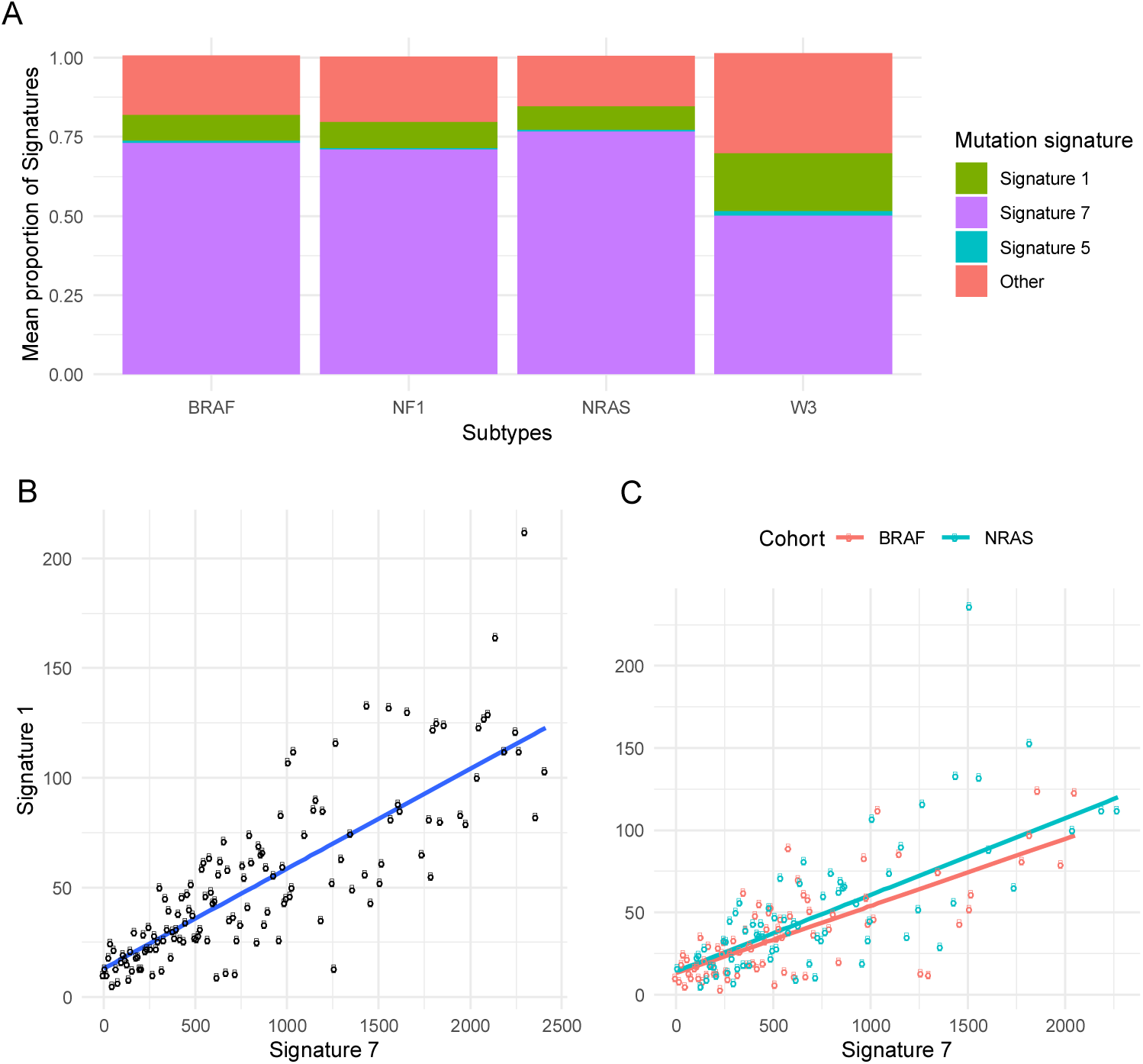
The Signature 7 UVR imprint predominates in melanoma and is tightly correlated to cell division Signature 1 mutations. **(A)** Mutation signature spectra and proportions in BRAF, NF1, NRAS, and W3 cutaneous melanomas. **(B)** Correlation analysis between somatic mutations due to extrinsic, UVR-driven Signature 7 mutations in cutaneous melanomas and intrinsic, clock-like Signature 1 mutations. **(C)** Correlation analysis between somatic mutations due to extrinsic, UVR-driven Signature 7 mutations in cutaneous melanomas subtypes and intrinsic, clock-like Signature 1 mutations (BRAF: red; NRAS: blue).

We then used Signature 1 and 7 to investigate the relationship between UVR damage and cell cycle, and show that cell division rate, predicted from Signature 1, tightly correlates with the total UVR-induced mutations in melanoma (Spearman ρ =0.82, *P*<10^−36^, Figure 3B). The correlation between Signatures 1 and 7 remains significant across all the subtypes (BRAF: Spearman ρ 0.70, *P*<10^−13^; NRAS: ρ 0.72, *P*<10^−12^; NF1: ρ 0.8, *P*<10^−8^; W3: ρ 0.64, *P*<10^−6^; Figure 3C). There is a marked difference in the increase of Signature 1 that is dependent on Signature 7 in both BRAF and NRAS melanomas (BRAF: robust regression with slope 0.033; *P*<10^−16^); NRAS: robust regression with slope 0.046; *P*<10^−12^). These data suggest UVR not only increases the total mutation burden by damaging DNA directly, but could also modify the total somatic TMB by affecting the dynamics of other mutational processes such as cell division. Importantly, these data show cell proliferation is coupled to UVR, not age, in BRAF melanomas.

### Ageing-associated mutations accumulate at different rates in males and females

Melanoma samples from male patients present an overall higher number of missense mutations than women in the TCGA SKCM cohort, adjusted for age and relevant clinical covariates(Gupta et al. 2015). We investigated the relationship between Signature 1 and sex, and critically, we observed that males present a higher number of Signature 1 mutations per age in the cohort overall (*P*<0.01, Mann-Whitney U test with Bonferroni correction; median male-to-female ratio 1.24; Figure 4A). The difference is still visible in the BRAF and NRAS subtypes (Figure 4B), although it is less statistically significant (*P*=0.1 for BRAF, *P*=0.6 for NRAS; Mann-Whitney U test with Bonferroni correction). For males, we found a significant rank-correlation with age (Spearman ρ 0.45, *P*<0.00023) but not for female samples (Spearman ρ 0.124, *P*=0.35; Figure 4A). Using multivariate negative binomial regression we found that sex affects the rate of mutation accumulation (*P*<10^−16^) and estimated the factor by which mutations increase per year in male and female samples (Supplementary Table 2; Figure 4C-4D). We observed only male samples presented a factor of mutation increase per year greater than one; and female melanomas showed no significant increase of mutations with age (Supplementary Table 2). The results remain robust when restricting the analysis to the molecular subtypes (except NF1 due to sample size), and show an increase in Signature 1 mutations with age only in males (Supplementary Table 2).

**Figure 4.**
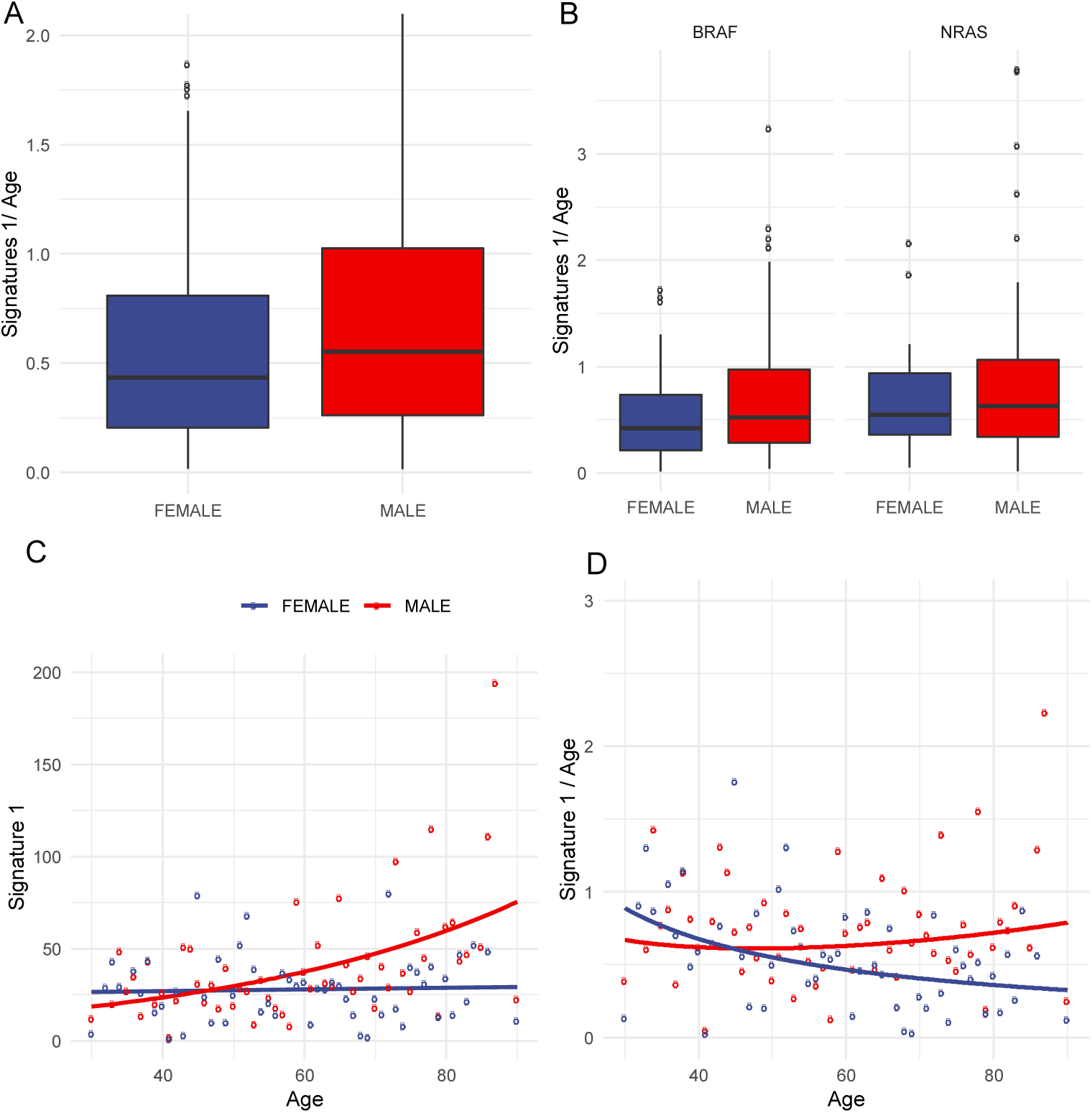
Melanoma in males accumulate more Signature 1 mutations. **(A)** Difference in Signature 1 mutations per age between males and females. **(B)** Difference in Signature 1 mutations per age between males and females by subtype (BRAF and NRAS). **(C)** Exponential model for accumulation of Signature 1 mutations by sex. This curve models the Poisson mean distribution of mutations at each age, with age-dependent rate. **(D)** Change of Signature 1 mutations per age over time, according to sex.

We next investigated whether this difference in the rate of mutation accumulation persists when accounting for the effect of UVR-driven Signature 7 mutations. Assuming that the number of Signature 1 mutations is proportional to the number of Signature 7 (see previous section Figure 3B, 3C), we investigated the ratios of Signature 1 to Signature 7 across melanomas, adjusted for the effect of Signature 7, and found that, when the effect of UVR on cell division is accounted for, there is little to no increase in Signature 1 mutations per year in either sex. Moreover, the multiplicative rate of increase of the ratio of Signature 1 to Signature 7 mutations per year turns out to be slightly smaller (by 0.01) in males (Supplementary Table 3) in all samples, across BRAF and NF1 subtypes, and is not detected in NRAS and W3. These data imply that the rate at which clock-like mutations accumulate per year depends on UVR, and this dependence is stronger in males than in females. In summary, our results suggest that the mutation burden due to cell division in melanoma is determined by UVR exposure, and males are more susceptible to UVR-induced cell proliferation. In contrast, females accumulate fewer Signature 1 mutations, at a slower rate, despite UVR exposure (Figure 5).

**Figure 5.**
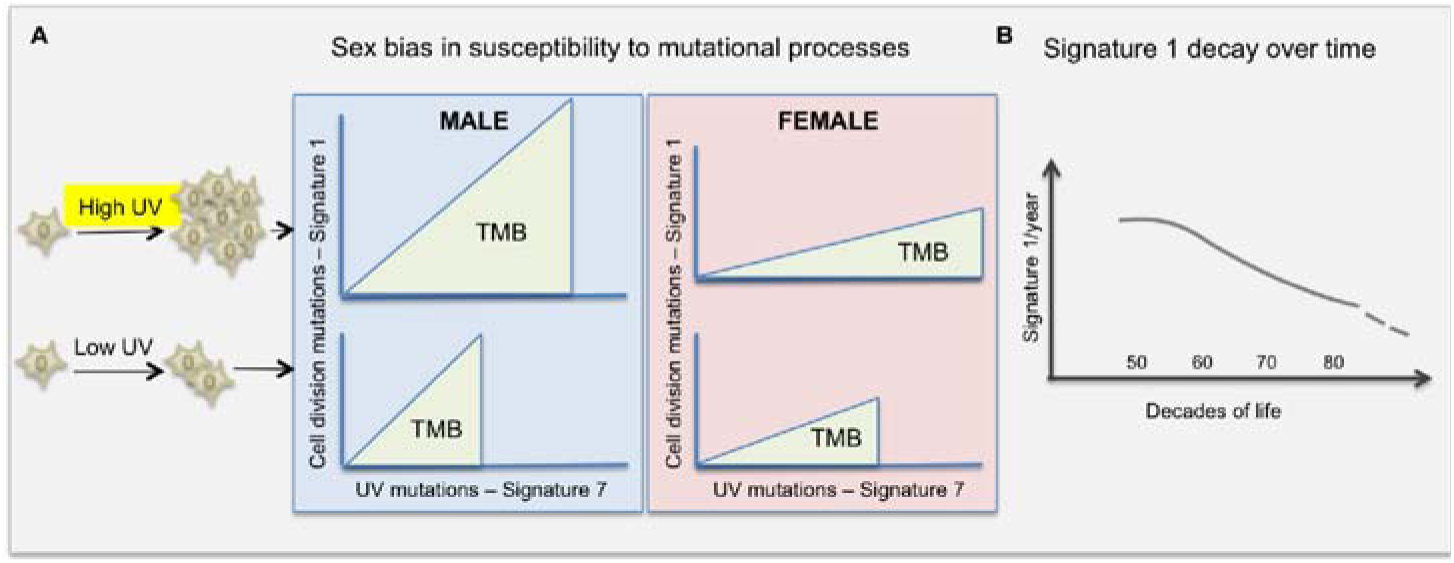
Summary findings. **A**. UV-light somatic mutations are strongly associated to the rate of cell-division mutations, suggesting that an extrinsic mutational process (UV) influences the intrinsic mutational process due to cell division. The rate of cell division in male melanoma is more strongly correlated to UV damage than in females. **B**. Cancer cells bear the genomic imprint of decreasing rate of cell division during ageing.

## Discussion

Sex and age differences have been observed across cancers and in melanoma(Gupta et al. 2015; Balch et al. 2001). Here we provide a novel mathematical framework to analyse, for the first time, the relationship between different damaging processes shaping the mutational landscape of human cancer; and how the mutational processes can reveal the effect of age and sex on carcinogenesis. We use the predominant mutational processes of the exomes of cutaneous melanomas to develop and test the new approach. Cutaneous melanoma DNA is imprinted primarily by (1) the clock-like changes due to cell division and (2) UVR-driven mutations. We reveal both processes increase during ageing and are tightly correlated, which poses the intriguing possibility that UVR exposure not only drives melanoma by damaging DNA directly, but also likely influences the intrinsic biological process of cell division and damage repair.

Using mathematical models allows us to compare the correlation of both signatures to age across the subtypes of melanoma. We observe the correlation of mutations to age is absent in BRAF melanomas, in sharp contrast to NRAS and NF1 melanomas, where we observe a gradual, long-term UVR exposure and cell division mutation burden increase as patients’ age. These data strongly support melanoma arises through different pathogenic pathways in the distinct molecular subtypes. The punctuated rate of mutation accumulation in BRAF melanomas could be driven by episodic acute sunburn, potentially uncoupling the correlation between age and mutation accumulation observed in other subtypes. Our tools and analysis reveal that the interaction between somatic mutations and environmental exposure is an important factor to consider in understanding tumour development.

Recent models examining the correlation between lifetime risk of a cancer and cell division show that tissues with higher cell turnover present an increased cancer incidence in the population(Tomasetti et al. 2016; Wu et al. 2016), which suggests tissues with high cell turnover require less environmental damage to drive tumourogenesis. Our results challenge this assumption in melanoma and suggest it is extrinsic processes such as UVR that modulate the contribution of cell division to mutation burden.

We show the natural decline in cell division as we age observed in healthy tissues(Wang and Hekimi 2015) is discernible in the genomic imprint of cancer cells. Recent *in vivo* work shows the cell division rate in stem cells declines with age(Tomasetti et al. 2019), and we validate these findings mathematically, for the first time in cancer cells, showing the rate of proliferative decline is not uniform across all melanoma subtypes. Our framework can be used to test if the age-driven decline in cell division varies depending on the tissue of origin, and whether the lineage-specific cell division decline mirrors a decrease in cancer incidence observed in the very elderly population.

Finally, we reveal a striking increase in the rate of cell division-linked mutations in males. This increase could be due to either an increase in the inherent proliferation rate of male melanocytes, or a decrease in the mutational repair of this mutational process as a consequence of extrinsic UVR-related mutations. A recent pan-cancer analysis has shown sex biases in mutational load, tumour evolution, mutational processes and at the gene level(Li et al. 2019); and our study suggests that sex differences in the melanoma TMB cannot be explained by lifestyle or age alone, and are likely a reflection of sex-specific biology and tumour evolution. Intriguingly, Signature 1 is increased in females in an age-adjusted, pan-cancer whole genome analysis(Li et al. 2019), whilst our results reveal a contrary sex bias in Signature 1 in melanoma. Overall our study provides new mathematical tools to examine the differences in the rate of mutation accumulation in cancer, and the dependencies of the mutational processes. We reveal a dependency of the intrinsic mutational process linked to cell division on environmental exposure in melanoma, and provide tools to examine how sex and age contribute to tumour development. Tumour burden, age and sex are variables known to influence the response to novel immunotherapies(Kugel et al. 2018; Samstein et al.) and future studies should address how these data can be leveraged to predict response to therapy and design strategies to improve overall survival.

## Materials and Methods

### Mutation data

The primary data are the somatic mutation calls from the The Cancer Genome Atlas (TCGA) MAF (skcm_clean_pairs.aggregated.capture.tcga.uuid.somatic.maf) of the whole-exome sequences of tumours from the TCGA skin cutaneous melanoma (SKCM) cohort(Akbani et al. 2015).

For each cutaneous melanoma sample we added information on whether the sample had mutations in ^V600^BRAF, ^G12/G13/Q61^NRAS, NF1 or none of these genes. (Five) samples had both ^V600^BRAF and ^G12/G13/Q61^NRAS mutations. These samples are likely to have acquired the NRAS mutations as a consequence of targeted therapy, and were excluded from the study. There were 24 (50) samples with mutations in both NF1 and ^V600^BRAF or ^G12/G13/Q61^NRAS, and they were classified as either BRAF or NRAS. Overall, 202 (163) samples have BRAF mutations, 109 (88) samples NRAS mutations, 52 (21) samples NF1 mutations, and the remaining 100 (51) samples were classified as triple wild type (W3). 278 of the samples had as primary site the trunk, head and neck, or extremities, while the remaining samples where from regional lymph nodes, distant metastases, or of unknown origin.

### Mutation signatures

For each sample and for each of K=96 types of single nucleotide substitutions (SNVs) in trinucleotide context, we count the number of times this substitution occurs relative to the abundance of the given context in the exome. The mutation catalogue is an integer vector counting the number of times each substitution out of a predefined alphabet appears in a sample, and such a catalogue can be approximated by a non-negative linear combination of mutation signatures,

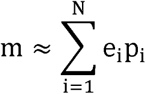

where the *p*_*i*_ ∈ ℝ^K^ are discrete probability distributions on the alphabet, called mutation signatures. The coefficient e_i_is called the exposure of the sample to signature p_i_. Since the entries of each signature add up to 1, the exposure of a signature measures the total number of mutations that can be blamed on the corresponding signature.

A set of 30 mutation signatures has been extracted and catalogued in the COSMIC database(Alexandrov et al. 2013). Of these signatures, signatures 1, 5 and 7 have been found to be relevant to melanoma(Cancer Genome Atlas Network. Electronic address and Cancer Genome Atlas 2015; Alexandrov et al. 2013). Signature 7 is known to be related to UV exposure. Signature 1 is dominated by C>T substitutions in NpG dinucleotide context, which is believed to be related to the spontaneous deanimation of 5-methylcytosine, and has been observed to accumulate at a constant rate over time in melanoma and some other cancers(Blokzijl et al. 2016; Alexandrov et al. 2015). We exclude signature 5 because ASC studies using liver organoids suggest this is an ageing mechanism independent of cell proliferation rate(Blokzijl et al. 2016).

We estimated the exposure to signatures 1 and 7 using two different methods. We first used the mutation signatures from the COSMIC database and determined the exposure of signatures relevant to melanoma using (R package deconstructSigs(Rosenthal et al. 2016)) We validated the approach by re-deriving the mutation signatures using a hierarchical Dirichlet process (R package hdp), taking into account the different molecular subtypes (BRAF, NRAS, NF1 and triple wild).

### Mathematical model of mutation accumulation

The number of mutations present at any given age can be described using a Poisson process(Podolsky et al. 2016) with time-varying mean λ(t). As suggested in(Podolsky et al. 2016), we use an exponential model λ(t) = N_0_e^αt^ for the mean. To estimate the effect of the different subtypes on the ratio of mutations by age we modelled the accumulation of mutations using a homogeneous Poisson process with age as offset,

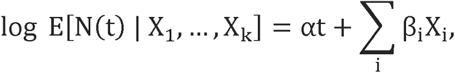

where the X_i_are the covariates (gender, site, subtype in our case). Note that in this model, the effect of each covariate on the expected ratio N(t)/t is multiplicative. Since the distribution of the mutation count data was found to be overdispersed (R package AES), we estimated a negative binomial regression instead of a Poisson regression.

While the difference in mutation ratios was found to be highly significant, the precise parameter values depend on the particular model used. In fact, the densities of the mutation frequencies at a given age range resemble a mixture of Poisson distributions, with one dominant component and smaller clusters at higher mutation counts. By fitting a Poisson mixture model, we found similar ratios between the coefficients associated to the different subtypes.

### Change in accumulation rate with age

Using the exponential model for mutation accumulation N(t) = N_0_e^αt^, the derivative of f(t) = e^αt^/t by t is 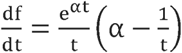. This expression is negative if α < 1/t, meaning a decrease of the ratio f(t) with t, and positive if α > 1/t, meaning an increase of f(t) with t. In particular, values of α < 0.01 would imply that for all ages t<100 we ratio of mutations by time is decreasing, while smaller ratios

### Ratio of Signature 1 to Signature 7 mutations adjusting for the effect of Signature 7

Using the exponential accumulation model, *N*(*t*) = *E N*_0_*e*^*αt*^, where E (for Extrinsic mutations) denotes the number of Signature 7 mutations (Poisson regression with the logarithm of Signature 7 as offset), we estimated the ratio *N*(*t*)/*E* of intrinsic, clock-like mutations when factoring out the extrinsic, Signature 7 mutations (Supplementary Table 3).

## Authors’ contributions

AV conceived the project. ML led the mathematical models and SF led the bioinformatics. AV, ML and SF interpreted the data and wrote the manuscript.

---

**Supplementary Table 1.** Proportion of subtypes in TCGA data set, average age of diagnosis by subtype, mean number of Signature 1 mutations and mean number of total mutations in BRAF, NRAS, NF1 and W3 cutaneous melanomas

|  | BRAF | NRAS | NF1 | W3 |
| --- | --- | --- | --- | --- |
| All | 172 | 96 | 44 | 84 |
| Male | 105 | 61 | 29 | 51 |
| Female | 67 | 35 | 15 | 33 |
| Mean Age | 53 | 59 | 70 | 63 |
| Mean Mutations | 595 | 844 | 1708 | 586 |
| Mean Signature1 | 33 | 47 | 70 | 32 |

**Supplementary Table 2.** The yearly rate at which Signature 1 mutations accumulate by sex and molecular subtype.

| Multiplicative accumulation of Signature 1 by year ($e^{\alpha}$) | MALE | FEMALE |
| --- | --- | --- |
| Total | 1.018 ($P < 10^{-8}$) | 1.0037 ($P = 0.401$) |
| BRAF | 1.01 ($P < 0.044$) | 0.997 ($P = 0.38$) |
| NRAS | 1.02 ($P < 0.01$) | 1.007 ($P = 0.39$) |
| NF1 | 1.001 ($P = 0.91$) | 0.992 ($P = 0.68$) |
| W3 | 1.049 ($P < 1.5 \cdot 10^{-8}$) | 0.988 ($P = 0.28$) |

**Supplementary Table 3.** The yearly rate a which Signature 1 mutations increases relative to Signature 7 mutations by sex.

| Multiplicative accumulation of Signature 1 / Signature 7 per year ($e^{\alpha}$) | MALE | FEMALE |
| --- | --- | --- |
| Total | 0.995 ($P < 10^{-12}$) | 1.005 ($P < 1.2 \cdot 10^{-8}$) |
| BRAF | 1.0046 ($P < 10^{-5}$) | 1.0127 ($P < 10^{-16}$) |
| NRAS | 1 ($P = 0.95$) | 1.0016 ($P = 0.4$) |
| NF1 | 0.985 ($P < 10^{-16}$) | 0.996 ($P = 0.13$) |
| W3 | 0.9998 ($P = 0.9$) | 0.965 ($P < 10^{-16}$) |

